# Snapshots from the Catalytic Landscape of Chalcone Isomerase

**DOI:** 10.64898/2026.09.01.748576

**Authors:** Jason R. Burke, Matthew A. Rangel, Vanessa I. Vasquez Meza, Hailey N. Mims, Thomas O. Darko, Kelwyn Ixcoy, Alfredo Ruiz Rivera, Jason Zhang, Emma R. Wolf-Saxon, Chad C. Moorman

## Abstract

Chalcone isomerase (CHI) catalyzes the cyclization of 3-ring scaffolds of flavonoids, a class of plant-based natural products important for nutrition and disease prevention. A persistent question has been whether the enzyme uses dynamics to facilitate conformational rearrangements of substrates within the active site. To help resolve this question, CHI was crystallized with phloretin, a flexible substrate analogue that cannot undergo cyclization. The crystal structure possesses eight protein molecules per asymmetric unit, revealing different active site conformations that accommodate different bound conformers of phloretin. Together, the structural snapshots depict a series of coordinated, dynamic chemical interactions that lower barriers to substrate rearrangements approaching bond formation. Differential scanning fluorimetry combined with mutational analysis and enzyme kinetics further confirm that phloretin binds to the enzyme active site and that it acts as a competitive inhibitor of CHI. Together these findings answer outstanding questions about the flexibility and dynamics of CHI catalysis, information that may be useful for future biosynthetic design and enzyme engineering goals. Overall, this work supports a catalytic model in which the CHI enzyme operates as a dynamic ensemble of structures necessary to facilitate catalytic substrate rearrangements.

## 1 Introduction

Enzymes enable life by enhancing rates of natural chemical reactions through dynamic adaptations that lower transition state energies; often these occur on timescales correlating with catalytic cycles (Henzler-Wildman et al. 2007, Frasier et al. 2009, Bhabha et al. 2011, Klinman 2013, Corbella et al. 2023, Hrmova 2026). Chalcone isomerase (CHI) is the plant enzyme that catalyzes the cyclization of the universal 3-ring backbone comprising all flavonoid metabolites (Bendar et al. 1998, Jez et al. 2000, Jez et al. 2002a, Jez et al. 2002b). The emergence of enzyme function in the catalytically nascent, ancestral CHI-fold proteins may have occurred around 500 million years ago in the earliest terrestrial plants (Ngaki et al. 2012, Cheng et al. 2018, Yonekura-Sakakibara et al. 2019). This is associated with active site repositioning of an arginine and the rapid movement of its guanidinium group (Kaltenbach et al. 2018). The specific interactions facilitated by the catalytic arginine have been elucidated in an X-ray crystal structure of CHI bound to a product molecule; however, the static nature of the product-bound state has left open questions of how active site dynamics contribute to CHI catalysis (Burke et al. 2019).

A second question concerning CHI dynamics is whether conformational changes in the enzyme facilitate necessary substrate rearrangements. It has been known for some time that cyclization of substrate chalcones requires a molecular reshuffling – from the thermodynamically-favored, ground state *s-cis* diene conformer to the sterically-disfavored, yet cyclization productive, *s-trans* diene conformer (Hur et al. 2004, Ruiz-Pernía JJ et al. 2007) (**Figure 1**). Molecular dynamics simulations support a model in which CHI can bind a 2′-hydroxychalcone substrate, either as a *s-cis* diene conformer or a *s-trans* diene conformer, and facilitate rearrangements from *s-cis* diene to *s-trans* diene within the active site in order to accomplish cyclization (Hur et al. 2004). However, additional empirical evidence has been lacking and a separate molecular dynamics study found that CHI binds only to the cyclization productive *s-trans* diene conformer; meaning to accomplish catalysis it must bind the thermodynamically disfavored conformer directly from its solution conformational equilibria (Ruiz-Pernía JJ et al. 2007). Beyond findings from these molecular simulations, there has been no evidence further supporting either model of CHI catalysis. However, it is notable that questions surrounding CHI catalysis are broadly applicable to other enzyme systems, and these two models effectively bookend a larger conversation concerning how enzyme active sites may be considered: Are they static, pre-organized, exquisitely molded architectures, designed to fit one transition state? Or, are they dynamic structural ensembles that are less efficient but possess greater functional flexibility (Maria-Solano et al. 2018)?

**Figure 1.**
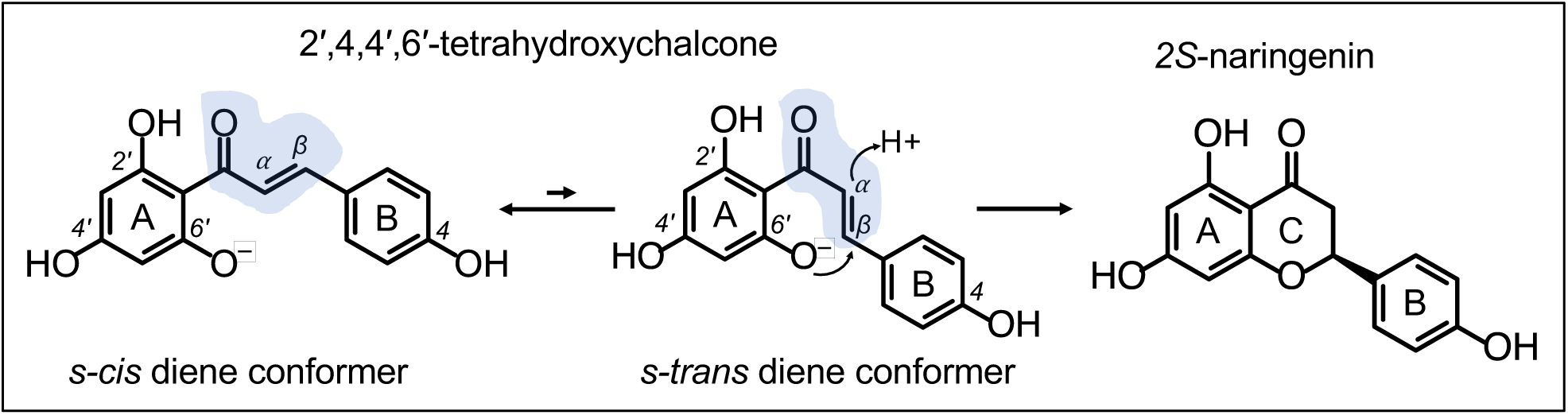
Conformational rearrangement and cyclization of 2′,4,4′,6′-tetrahydroxychalcone. Cyclization of 2′,4,4′,6′-tetrahydroxychalcone requires conformational rearrangement from a ground state, *s-cis* diene conformer to a sterically-disfavored, *s-trans* diene near attack conformer, which is cyclization productive. Existing dynamic models disagree on whether the CHI enzyme facilitates the *s-cis* diene to *s-trans* diene conformational rearrangement prior to CHI-catalyzed regioselective closing of the C ring, to form 2*S*-naringenin.

To help resolve outstanding questions concerning the dynamic catalytic capabilities of CHI, we here report a structure of a CHI from *Medicago truncatula* (Barrelclover), termed *Mt*CHI-I, bound to the substrate analogue phloretin. The *Mt*CHI-I enzyme was selected for this study because it is amenable to crystal ligand soaking and possesses 8 structurally diverse protein molecules per asymmetric unit in a solved crystal form (Burke et al. 2019). The substrate analogue phloretin was selected because it binds well to *Mt*CHI-I and possesses rotational freedom that can provide insight into how different possible substrate conformations are accommodated within the enzyme active site. The X-ray crystal structure presented here reveals *Mt*CHI-I enzyme active sites with eight different conformers of bound phloretin, each stabilized by transient and stable coordinating H-bonds from the catalytic arginine and other nearby amino acids. Protein melting experiments confirm that key active site amino acids, including the catalytic arginine, provide H-bond interactions that stabilize the *Mt*CHI-I-phloretin interaction. Enzyme kinetics experiments reveal that phloretin acts as a competitive enzyme inhibitor of the studied *Mt*CHI-I, and although phloretin is not produced in *Medicago truncatula*, this finding may have metabolic implications for other species in which phloretin and other reduced dihydrochalcones are made. Together, these results provide evidence for the importance of active site dynamics in catalytic cycles performed by chalcone isomerase enzymes. The resulting biochemical model may be useful in future efforts aimed at bioengineering and biocatalyst design with goals of generating novel flavonoids through active site flexibility and improved flavonoid production efficiency in host systems.

## 2 Results

### 2.1 Selection of Phloretin as a Substrate Analogue for Crystallography Studies

Two analogues of the CHI substrate, 2′,4,4′,6′-tetrahydroxychalcone, were examined for binding to *Mt*CHI-I. Isoliquiritigenin (2′,4,4′-trihydroxychalcone) is inert toward catalytic cyclization by *Mt*CHI-I and other type-I CHIs due to its lack of a 6′OH (Shimada et al. 2003, Ralston et al. 2005). Phloretin is a reduced dihydrochalcone that is inert toward cyclization because it lacks the alkene functional group (Gosch et al. 2009). Differential scanning fluorimetry (DSF) was used to measure enzyme binding of these two substrate analogues, as compared to the reaction product naringenin. Titration of isoliquiritigenin into *Mt*CHI-I generates a positive maximum thermal shift (ΔTm_max_ = 4.2 ± 0.3 °C), indicating a protein fold–stabilizing effect, and a single site binding equation was used to solve for the apparent equilibrium binding dissociation constant (app. K_d_) of 131 ± 68 μM (**Figure 2A**). Since protein-ligand binding measured via DSF does not account for the temperature dependence of binding (Simeonov 2013), apparent K_d_ values (app. K_d_) are used here. Titration of phloretin into *Mt*CHI-I generates a more significant binding response, with a ΔTm_max_ of 9.8 ± 1.0 °C and app. K_d_ of 26 ± 1 μM (**Figure 2B**). For comparison, titration of racemic, (*2S/2R*)-naringenin into *Mt*CHI-I generates a ΔTm_max_ of 4.1 ± 0.1 °C and app. K_d_ of 598 ± 161 μM (**Figure 2C**). Racemic, (*2S/2R*)-naringenin was used due to the difficulty of obtaining pure *2S*-naringenin. Together, these results identified phloretin as a relatively strong interactor and stabilizer of *Mt*CHI-I, and a promising substrate analogue to soak into preformed protein crystals of *Mt*CHI-I grown under previously established conditions (Burke et al. 2019).

**Figure 2.**
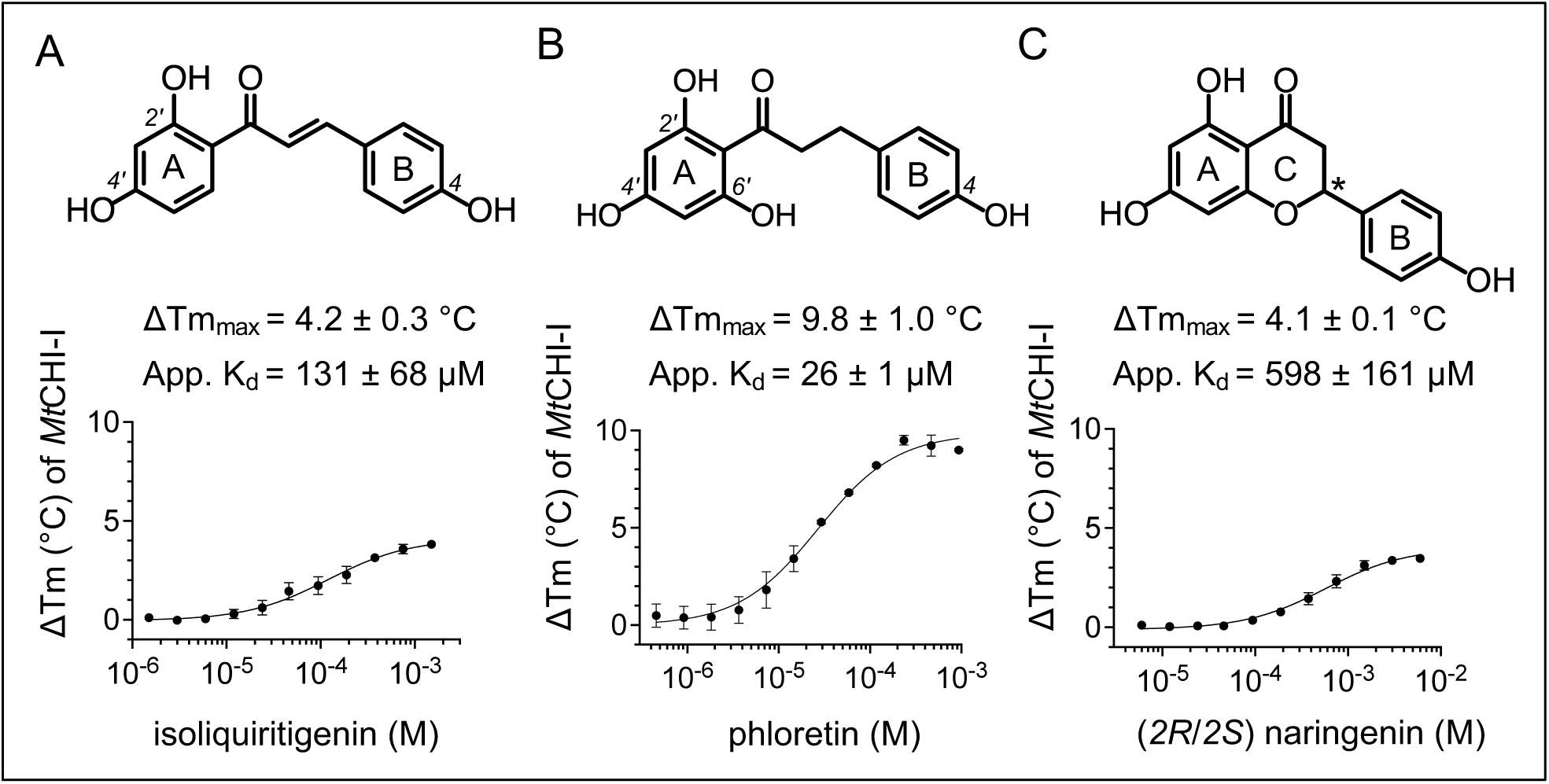
DSF derived parameters of *Mt*CHI-I binding to chalcone and flavonoid metabolites. **A)** The structure of isoliquiritigenin, ΔTm_max_, and app. K_d_ values, and dose-response data of isoliquiritigenin interactions with *Mt*CHI-I. **B)** The structure of phloretin, ΔTm_max_, and app. K_d_ values, and dose-response data of phloretin interactions with *Mt*CHI-I. **C)** The structure of racemic (*2R/2S*)-naringenin (racemic at the asterisks), ΔTm_max_, and app. K_d_ values, and dose-response data of (*2R/2S*)-naringenin interactions with *Mt*CHI-I. Error bars are standard deviations from the average of four to five independent experiments.

### 2.2 Crystallographic Insights into the Catalytic Plasticity of CHI

A previous protein X-ray crystal structure of *Mt*CHI-I bound to its product, 2*S*-naringenin, revealed 8 protein molecules in the asymmetric unit (Burke et al. 2019).

Based on this product-bound structure, we hypothesized that differences between the protein molecules in the asymmetric unit may reveal catalytically-important enzyme conformers which direct structural changes in substrates, to facilitate cyclization. With this in mind, phloretin was soaked into preformed apo *Mt*CHI-I crystals and the structure was solved (**Table 1**). In the *Mt*CHI-I-phloretin structure, each of the eight *Mt*CHI-I molecules within the asymmetric unit possess varying degrees of structural deviations relative each other, as evidenced by pairwise, C*a*-based structural alignments, with root mean square (RMS) values ranging from 0.168 Å to 0.596 Å (**Figure 3A, 3B).** A C*a*-based structural alignment and superposition of all eight *Mt*CHI-I molecules reveals the diversity of conformers sampled by the phloretin substrate analogues in coordination with the catalytic arginine (R37) within the active site (**Figure 3C**). A structural alignment of the A ring atoms of the eight phloretin molecules shows the range of phloretin conformer rotation about dihedral angle 2; the rotation required for the ring closure of substrate (Hur et al. 2004) (**Figure 3D**). For each *Mt*CHI-I molecule, a polder omit map supports the placement and refined conformation of the phloretin molecule in each active site (**Figure 3E)** (Liebschner et al. 2017).

**Figure 3.**
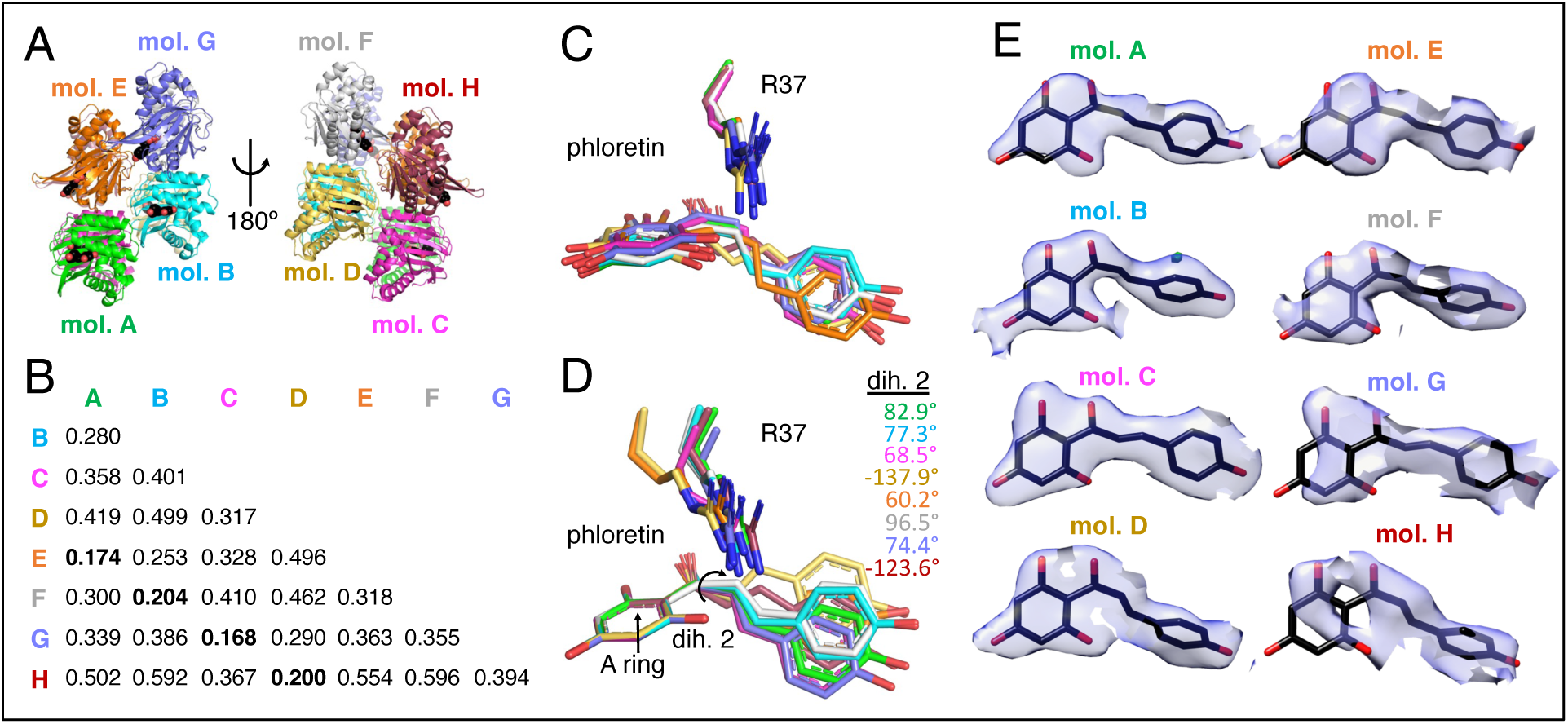
Molecular alignments from the *Mt*CHI-I-phloretin X-ray crystal structure. **A)** The asymmetric unit contains eight *Mt*CHI-I-phloretin molecules, shown as cartoon, with phloretin as space-filling in black and red. **B)** A matrix of RMS values from pairwise, C*a*-based structural alignments of *Mt*CHI-I molecules, labeled A-G. **C)** Superposition of C*a*-based structural alignments of all eight *Mt*CHI-I molecules, focused only on phloretin in coordination with the catalytic arginine (R37) **D)** Structural alignments based on the A ring atoms of the eight phloretin molecules reveals the range of phloretin conformer rotation about dihedral angle #2 (dih. 2). **E)** Polder omit maps supporting unbiased electron density of refined phloretin molecules in the enzyme active sites.

**Table 1.**
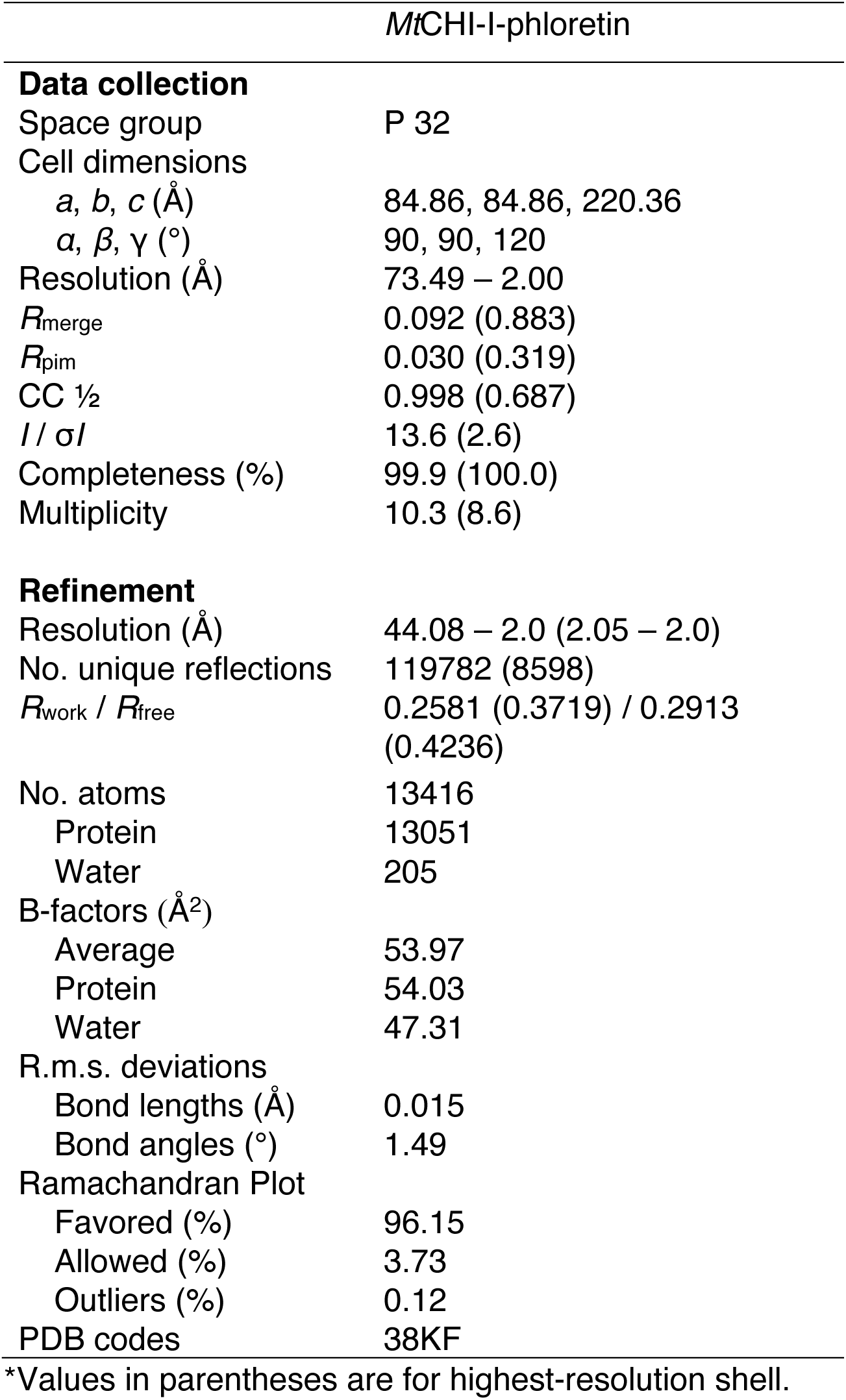
X-ray crystallography data collection and refinement statistics. Parentheses are for the highest resolution shell.

The eight *Mt*CHI-I-phloretin structures reveal a possible trajectory of the dynamic enzyme-substrate complexes leading to substrate cyclization. Furthest from cyclization is the conformer with the greatest interatomic distance between phloretin atoms (Ph)2′O and (Ph)C3, which are analogous to the ring-closing atoms in chalcone substrates. In the pose called “mol. D”, the phloretin (Ph)2′O-(Ph)C3 interatomic distance is 3.98 Å, revealing a relatively extended conformation (**Figure 4A**). “Mol. D” and “Mol. H” are therefore the phloretin conformations most similar to the nonproductive ground state conformer of 2′,4,4′6′-tetrahydroxychalcone, shown in **Figure 1**. In intermediate poses closer to cyclization, the phloretin (Ph)2′O-(Ph)C3 interatomic distance closes in – to a distance between 3.30 Å and 3.18 Å – and mol. F, C, A, E and G reveal how this is modulated by a few additional transient hydrogen bonds between the catalytic arginine (R37) and phloretin: specifically, (R37)Nɛ forms H-bond interactions with (Ph)2′O and (Ph)C2, as atoms of phloretin are positioned by the catalytic arginine to move closer to each other (**Figure 4A**). The phloretin conformation that best approximates the near attack, cyclization-productive conformer occurs in mol. B, which measures the smallest interatomic distance between (Ph)2′O-(Ph)C3, of 3.09 Å (**Figure 4A**). Here, three critical H-bond interactions are provided by the guanidinium of R37 when they are at their closest to phloretin, underscoring the importance of these interactions for catalysis. Specifically, (R37)N_ƞ2_ aligns with (Ph)2′O and is positioned to stabilize the anionic electrostatic state on the oxygen, which is necessary to direct the ring closure of chalcone substrates (Jez et al. 2000). At the same time, (R37)Nɛ is positioned to direct asymmetric proton transfer from the guanidinium Nɛ atom onto (Ph)C2, which was previously measured by isotope labeling and NMR (Burke et al. 2019). This step, which consists of direct proton transfer from (R37)Nɛ to Cβ of 2′,4,4′,6′-tetrahydroxychalcone (analogous to (Ph)C2 of phloretin), is the rate-limiting step the reaction (Ruiz-Pernía JJ et al. 2006, Ruiz-Pernía JJ et al. 2008). Together, these structural observations made from the conformational ensemble paint a dynamic portrait of CHI during catalysis: CHI enzymes are equipped to bind chalcone substrates as ground state, *s-cis* diene conformers, followed by enzyme-mediated rearrangements, which stabilize barriers of substrate rotations, to near attack, *s-trans* conformers ready for cyclization.

**Figure 4.**
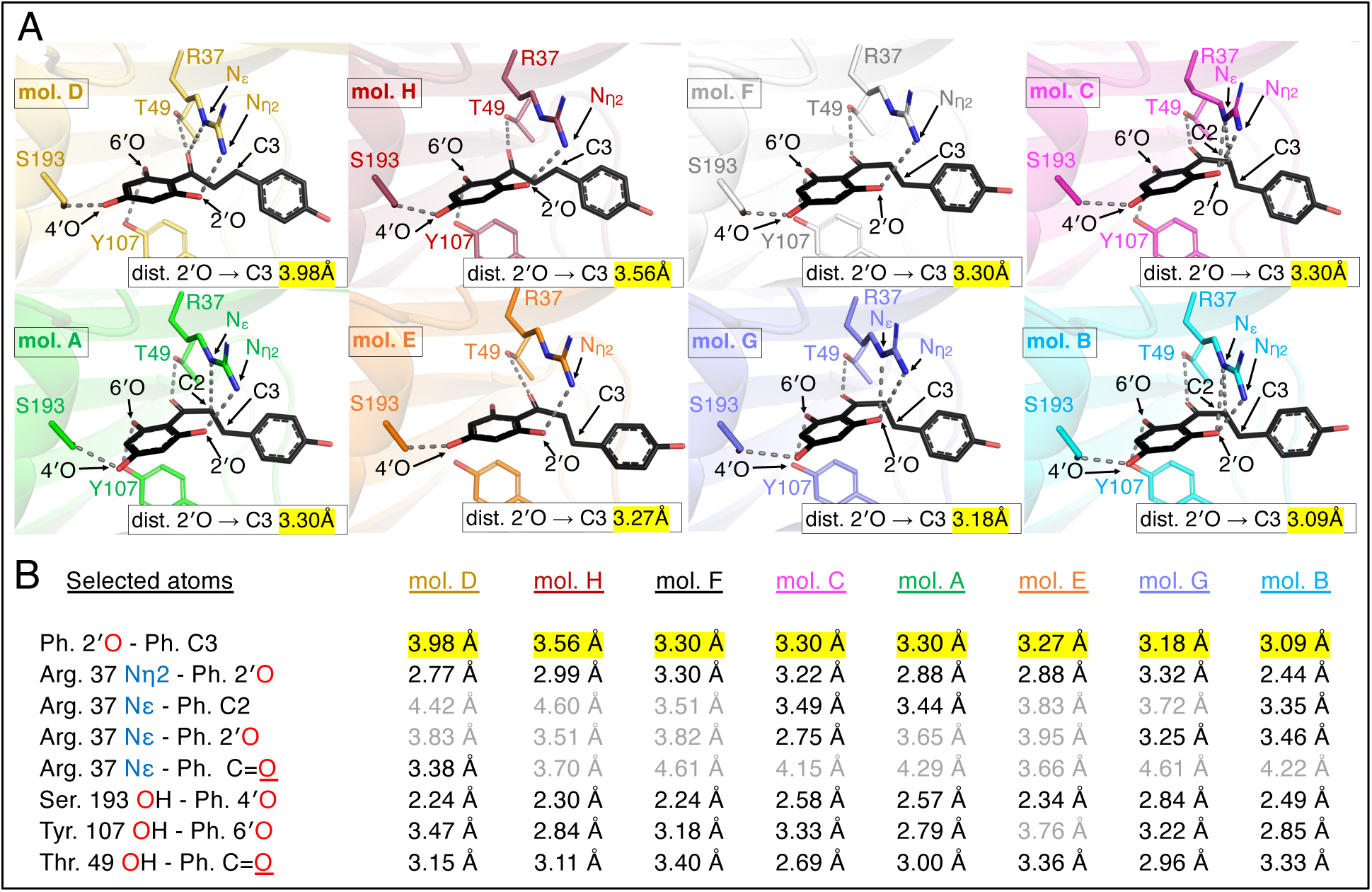
Crystallographic complex poses and measured distances in the *Mt*CHI-I-phloretin X-ray structure. **A)** Crystallographic poses of phloretin bound within the active sites of *Mt*CHI-I molecules. Differences in structures reveal how active site dynamics stabilize the reorientation of phloretin from a near ground state conformer (mol. D, top left) to a near attack conformer (mol. B, bottom right), as measured by the shrinking distances between the atoms analogous to the ring closing atoms of chalcone: (Ph)2′O and (Ph)C3. **B)** Interatomic distances measured between key H-bond atoms of *Mt*CHI-I mols. A-H and phloretin. Distances less than 3.5Å are shown in black, distances greater than 3.5Å are shown in gray.

Across the observed crystallographic *Mt*CHI-I-phloretin poses, persistent H-bond stabilization interactions occur, but with fluctuating H-bond distances (**Figure 4B**). The most critical of these may be the enzyme-substrate H-bond that occurs between (R37)N_ƞ2_ and (Ph)2′O, which is maintained through all observed poses. This interaction orients and activates the ring-closing oxygen of the substrate, and the bond distance is closest in the final pose, mol B (**Figure 4A**). In addition, other measured H-bond distances reveal that the active site position and orientation of the A ring of phloretin is very consistently maintained by three amino acid sidechains: threonine 49 (T49); serine 193 (S193), and tyrosine 107 (Y107) (**Figure 4B**).

### 2.3 Solution Studies of Phloretin Binding within the *Mt*CHI-I Active Site

To validate the observed crystallographic H-bond interactions between phloretin and the active site residues of *Mt*CHI-I, the following variants were purified: Y107F, S193A, R37K and T49A. Each of these variants is designed to abolish key *Mt*CHI-I-phloretin H-bond interactions observed in the crystal structure (**Figure 4A).** Previously, we showed titration of phloretin into wild type *Mt*CHI-I generates a maximum thermal shift, ΔTm_max_, of 9.8 ± 1.0 °C and produces a binding interaction with an apparent K_d_ of 26 ± 1 μM (**Figure 2B**). This result is republished in **Figure 5** to establish a basis for comparison with variant versions of the protein. In testing these variants, we find: T49A generates a ΔTm_max_ of 4.0 ± 1.1 °C and app. K_d_ of 59 ± 14 μM; R37K generates a ΔTm_max_ of 6.8 ± 0.3 °C and app. K_d_ of 46 ± 6 μM; S193A generates a ΔTm_max_ of 7.4 ± 0.2 °C and app. K_d_ of 34 ± 3 μM, and Y107F generates a ΔTm_max_ of 10.2 ± 1.1 °C and app. K_d_ of 25 ± 8 μM (**Figure 5**). These results confirm that the H-bonds between phloretin and the sidechain atoms of Ser193, Arg37 and Thr49, observed in the *Mt*CHI-I-phloretin crystal structure, are also important for solution binding. On the other hand, the DSF results for the variant Y107F are very similar to wild type, indicating that the H-bond observed between the Tyr107 sidechain hydroxyl and the 6′-OH of phloretin may not play a significant role in phloretin binding. Other than Y107F, these results align closely with enzyme kinetics experiments which show that each of these amino acid substitutions have demonstrated large reductions in the catalytic efficiency of *Mt*CHI-I toward 2′,4,4′,6′-tetrahydroxychalcone (Burke et al. 2019).

**Figure 5.**
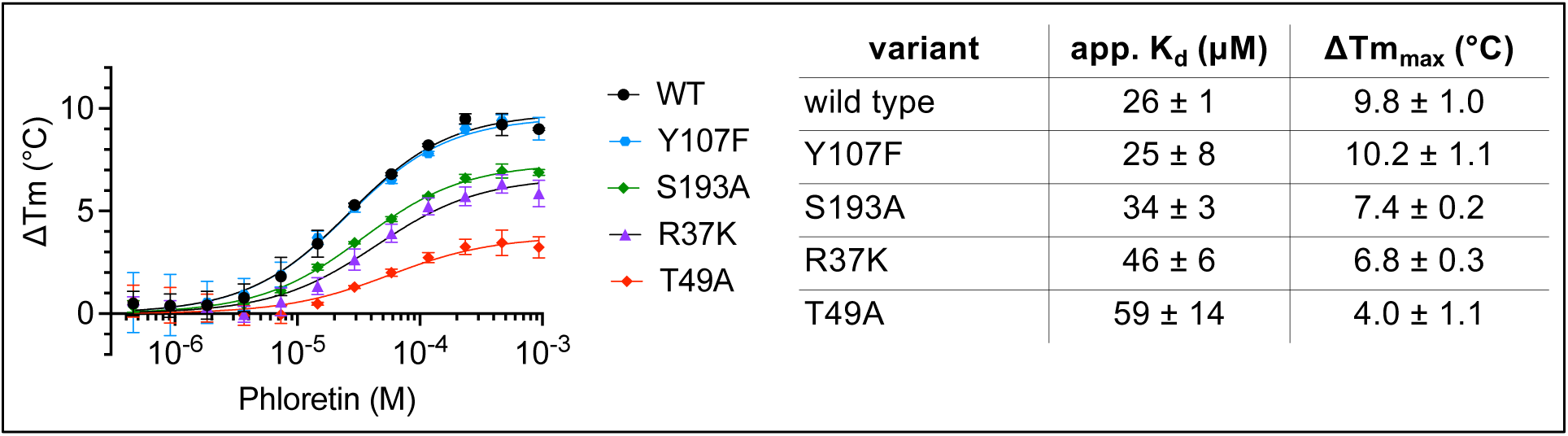
Solution studies of *Mt*CHI-I-phloretin binding and the effect of mutations to observed active site interactions. DSF was used to measure T_m_ values for *Mt*CHI-I at multiple phloretin concentrations for wild type *Mt*CHI-I, Y107F, S193A, R37K and T49A. The values for app. K_d_ and ΔTm_max_ were fit using a single-site binding equation. As measured by DSF, phloretin interactions with *Mt*CHI-I variants T49A, S193A and R37K, result in changes to ΔTm_max_ and app. K_d_ that are consistent with weaker binding, while Y107F more closely matches wild type. Error bar values are standard deviations from the mean of four or more independent experiments.

### 2.4 Phloretin is a Competitive Enzyme Inhibitor of *Mt*CHI-I

The orientation and positioning of phloretin binding within the *Mt*CHI-I active site suggest that phloretin may act as a competitive enzyme inhibitor. Steady state enzyme kinetics were performed using 2′,4,4′,6′-tetrahydroxychalcone as a substrate in the presence of varying concentrations of phloretin. A double reciprocal Lineweaver-Burk plot confirms that phloretin inhibits *Mt*CHI-I through a competitive mechanism, as greater phloretin concentrations increase apparent K_m_ (as observed by the decreasing magnitude of negative values at the X-intercept), but have no effect on V_max_ (observed at the Y-intercept) (**Figure 6**). From the enzyme inhibition experiments, the inhibitory constant, K_i_ value, averaged from the three inhibitor concentrations, is calculated to be 32 ± 4 μM. This is in agreement with the app. K_d_ of 26 ± 1 μM from the phloretin *Mt*CHI-I binding experiment, as measured by DSF (**Figure 2B**).

**Figure 6.**
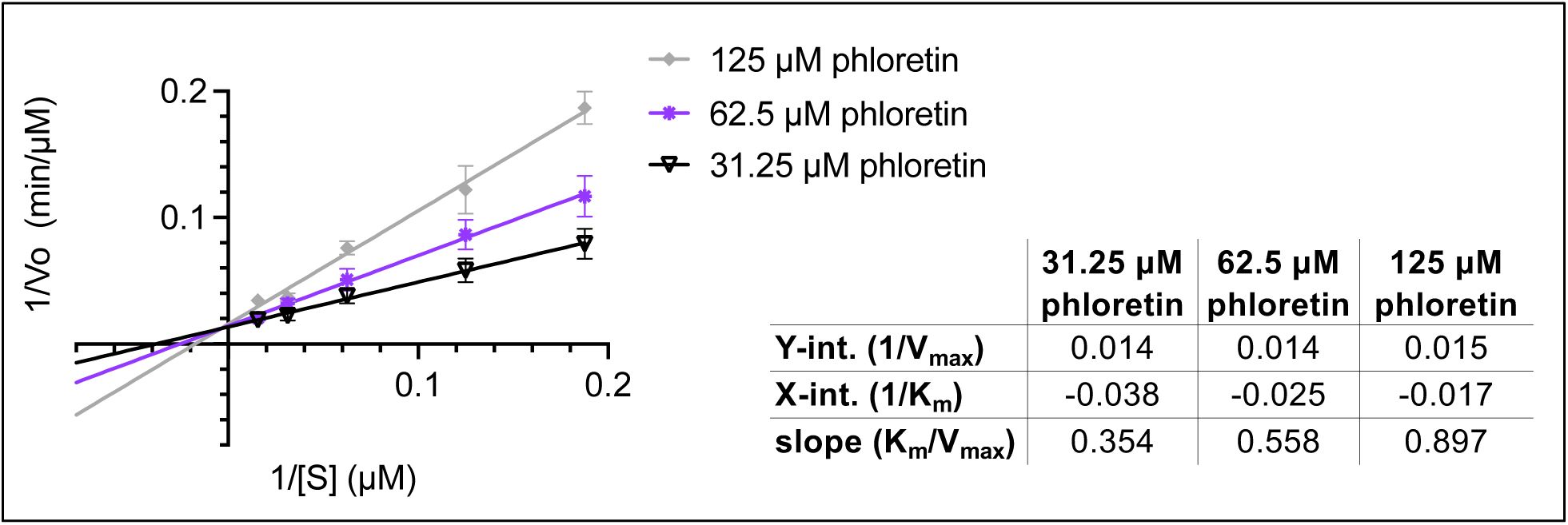
A double-reciprocal Lineweaver-Burk plot of *Mt*CHI-I enzymatic activity in the presence of phloretin is indicative of the type of enzyme inhibition caused by phloretin. As the inhibitor concentration increases, the V_max_ is unchanged (Y-intercept) while the K_m_ increases (X-intercept), suggesting phloretin acts through a competitive enzyme inhibition mechanism. Error bar values are standard deviations of three to four independent experiments.

## 3 Discussion

The findings presented here answer outstanding questions about the enzyme mechanism of CHI by revealing structural dynamics between the active site and substrate. The data reveal how phloretin, a substrate analogue and competitive inhibitor of CHI, binds within the active site of *Mt*CHI-I and exhibits conformations similar to the substrate in a non-productive ground state, as well as intermediate poses leading to cyclization. Together, this array of crystallographic poses provides new experimental support for a mechanistic model initially proposed by Hur and colleagues in which the CHI active site facilitates dynamic substrate rearrangements in order to efficiently drive catalysis (Hur et al. 2004). This study also helps expand on previous findings which sought to explore CHI dynamics in the context of protein evolution and showed that catalytic gains in CHI are most firmly associated with active site repositioning of the catalytic arginine and rapid movement of its guanidinium group on the nanosecond timescale (Kaltenbach et al. 2018). In the present work, we see how the arginine’s guanidinium movements facilitate changing H-bond interactions that stabilize substrate rearrangements throughout steps in the catalytic cycle.

One notable limitation of the present work is that previous models of CHI, such as the one used by Hur and colleagues, were based upon evidence from a Type-2 CHI, while the present work examines only a Type-1 CHI (Hur et al. 2004). Type-2 CHIs, which are generally restricted to the legume family (Fabaceae), have the evolved capability to bind water molecules in the back of the active site, enabling cyclization of 2′-deoxychalcone substrates which feed into the isoflavone pathway (Ralston et al. 2005, Burke et al. 2019). Crystal structures of a Type-2 CHI from *Medicago sativa* display the active site arginine in a position that is bent out of the active site; a conformation for which the significance is not well understood and which initially led to models wherein the arginine was not important for catalysis (Jez et al. 2000, Jez et al. 2002, Hur et al. 2004, Ruiz-Pernía JJ et al. 2007). However, recent work has revealed that the active site arginine is essential for catalysis in both Type-I and Type-2 CHIs, despite the structural differences between the arginine position in the different structures (Burke et al. 2019).

The flavonoid pathway of plants produces thousands of natural product molecules, many of which are investigated for their metabolic and tissue-specific roles in human health (Williamson et al. 2018). Given the molecular diversity and intrinsic difficulty of synthesizing and purifying key flavonoid natural products, interest in bioengineering and biosynthesizing flavonoid production, through the use of CHI-fold proteins and related pathway enzymes, is growing (Nabavi et al. 2018). However, within this effort there has not yet been any investigation of the structural basis for CHI catalytic promiscuity, nor the potential for lab-based CHI evolvability as it relates to engineering novel flavonoids. In nature, CHI-fold proteins are tunable to various functions which suggests they may serve as adaptable starting points for modern biosynthetic enzyme engineering goals. For example, although the most ancient CHI-fold proteins may have been nonenzymatic and only possessed fatty-acid binding functions in primary metabolism (Ngaki et al. 2012), extant CHI enzymes approach diffusion-limited catalysis while also encompassing family-specific (Type-1 *vs.* Type-2) catalytic functions (Jez et al. 2000, Jez et al. 2002b, Shimada et al. 2003, Dastmalchi et al. 2015, Kaltenbach et al. 2018, Burke et al. 2019, Wang et al. 2022). Chalcone Isomerase-like (CHIL) proteins, which share the CHI-fold yet lack CHI enzymatic activity, may be even more remarkable for their varied capabilities by which they enhance flavonoid production, through binding to chalcone synthase (CHS) (Morita et al. 2014, Jiang et al. 2015, Waki et al. 2020, Ni et al. 2020, Xu et al. 2022, Wolf-Saxon et al. 2023, Lewis et al. 2024). Functionally, the CHIL-CHS complex down regulates production of the side product, coumaroyltriacetic acid lactone (CTAL), and upregulates flavonoid synthesis via 2′,4,4′,6′-tetrahydroxychalcone production (Waki et al. 2020, Ni et al. 2020). This further enhances naringenin production and partitioning toward isoflavonoids in biotransformed yeast (Xu et al. 2022, Raytek et al. 2025). The CHIL-CHS complex, which may be best structural evidence of a flavonoid metabolon to emerge thus far, reveals atomic details of how CHIL reshapes the CHS active site to modulate product specificity (Imaizumi et al. 2026, Wang et al. 2026). Finally, fungal CHI-fold proteins in *Saccharomyces cerevisiae* lack any catalytic activity or fatty acid–binding capabilities, but instead bind heme proteins, further expanding the diverse functions of the CHI-fold (Schmitz et al. 2023). Taken together, better structural models of CHI-fold enzymes, along with other flavonoid producing enzymes and regulatory complexes, will enhance broader structure-based dynamic models of catalysis important for computational and rational design of novel biocatalysts (Crean et al. 2020).

Phloretin is a dihydrochalcone natural product most notably found in *Malus domestica* Borkh. (apple), where it modulates growth and protects against herbivory, among other functions (Dare et al 2013, Dare et al. 2020). Phloretin is biosynthesized from *p*-coumaroyl-CoA through a shunt in the flavonoid biosynthesis pathway that utilizes naringenin chalcone reductases (NCRs) to form reduced chalcones (Gosch et al. 2009, Ibdah et al. 2014, Yauk et al. 2024). As such, the formation of dihydrochalcone diverts CHI substrates and interferes with the formation of flavonoids (Gosch et al. 2009, Dare et al. 2020). Here we show that phloretin also directly inhibits plant CHI by acting as a competitive enzyme inhibitor of *Mt*CHI-I *in vitro*, with a K_i_ of 32 ± 4 μM. This suggests phloretin may also negatively regulate CHI activity *in vivo* through a feedback mechanism favoring dihydrochalcone production over production of naringenin in *Malus domestica* Borkh. While it is notable that *Medicago truncatula* does not produce phloretin, phloretin inhibition of *Mt*CHI-I suggests that phloretin may in fact be a relatively promiscuous inhibitor of CHI enzymes. For example, it has been shown that phloretin directly inhibits a gut microbiome CHI which metabolizes flavanones in *Eubacterium ramulus* (Herles et al. 2004); despite the fact that this enzyme does not possess a CHI-fold and is not evolutionarily related to plant CHI (Thomsen et al. 2015).

In summary, we here provide the first structural evidence of the process of chalcone ring closure within the CHI active site, as directly facilitated by coordinated movements of the substrate and a catalytic arginine. In doing so, this work provides an updated model supporting prior studies of CHI dynamics-enabled catalysis that will be useful for future biocatalytic design goals.

## 4 Methods

### 4.1 Protein Expression and Purification

A pET-21b protein expression plasmid containing the gene for *Medicago truncatula*, *Mt*CHI-I (UniProt: B7FJK3), was transformed into BL21 DE3-pRIL cells (Agilent Technologies). Cells were grown in Terrific Broth (Thermo Fisher Scientific) and induced overnight at 20°C. Cells were lysed in a buffer containing 150 mM NaCl, 1 mM 2-mercaptoethanol and 25 mM Tris-HCl (pH 8.0), and His-tagged proteins were purified by Ni-affinity chromatography using HisPur Ni-NTA resin (Thermo Fisher Scientific). The His tag was cleaved at 4°C by bovine thrombin (EMD Millipore Corp.). After overnight dialysis at 4°C, the CHI proteins were repurified over a Ni-affinity column by collecting the flow-through. Protein concentration and purity was measured on a NanoDrop 2000 (Thermo Fisher Scientific). Genetic variants of *Mt*CHI-I (R37K, T49A, Y107F, S193A) had been generated in a previous study (Burke et al. 2019). These proteins were expressed and repurified as described above.

### 4.2 Differential Scanning Fluorimetry

Assays were performed using SYPRO orange dye (1:625 dilution) and 6 µM *Mt*CHI-I protein against dilution series of: racemic (2*R*/2*S*)-naringenin (Sigma Aldrich); isoliquiritigenin (4,2′,4′-trihydroxychalcone, Indofine); and phloretin (TCI). Reactions were run in 20 µl reaction volumes and in a buffer containing 150 mM NaCl, 25 mM Tris pH 8.0, 2 mM DTT and 2% DMSO. Due to low solubility, 2% DMSO was used for all reactions to enhance ligand solubility. Experiments were performed on a QuantStudio 3 Real-Time PCR (Thermo Fisher Scientific) and analyzed using Protein ThermalShift software (Thermo Fisher Scientific). Replicates of 3 or more experiments were conducted for each condition and Prism 10.0 software (GraphPad) was used to fit a single site binding equation to the data to fit ΔTm_max_ and app. K_d_ values.

### 4.3 Protein Crystallography, Structure Determination and Analysis

For protein crystallography, *Mt*CHI-I was first purified by Ni-affinity chromatography as described above, then further purified by FPLC on a resource Q anion exchange column (Cytiva), and finally by gel-filtration chromatography on a Superdex 200 column (Cytiva) with an eluent consisting of 200 mM NaCl, 2 mM DTT and 25 mM Tris-HCl (pH 8.0). *Mt*CHI-I crystallized by hanging drop vapor diffusion at 30 mg/ml and under conditions previously described (Burke et al. 2019). For the generation and characterization of *Mt*CHI-I-phloretin complexes, *Mt*CHI-I crystals were harvested, soaked twice for several hours in a solution consisting of 30% (v/v) PEG1K and 100 mM Tris pH 8.5. Finally, the *Mt*CHI-I crystals were soaked for one week in the reservoir solution also containing 20 mM phloretin. Crystals were flash frozen in liquid N_2_ without additional cryoprotectant and X-ray diffraction data were collected at 100 K the Advanced Light Source beamlines 8.2.1 and 8.2.2, Lawrence Berkeley National Laboratory. Data were indexed and integrated with iMOSFLM (Battye et al. 2011), scaled with AIMLESS (Evans et al. 2011), solved by molecular replacement in space group P3_2_ using the program PHASER (McCoy et al. 2007), and using a search model derived from the structure of *Arabidopsis thaliana* CHI (PDB:4DOI) (Ngaki et al. 2012). Models were built in Coot (Emsely et al. 2010) and refined in PHENIX with ligand restraints for phloretin generated from eLBOW (Adams et al. 2010). Additional steps for refining the *Mt*CHI-I crystals, which showed combined effects of merohedral twinning and rotational pseudosymmetry, were used by applying a twin operator [h,-h-k,-l], and have previously been described in detail (Burke et al. 2019). Polder omit maps were constructed in PHENIX to assess unbiased ligand density (Liebschner et al. 2017), and all pairwise C*a*-based structural alignments were performed in PyMOL (Schrodinger, LLC). Statistics for data collection/processing and model refinement are shown in **Table 1**. Specific PDB information can be found at the Protein Data Bank for entry 38KF, Extended PDB ID pdb_000038KF.

### 4.4 Enzyme Kinetics

All kinetic measurements are based on a decrease in substrate absorbance of 2′,4,4′,6′-tetrahydroxychalcone, at lambda max = 390 nm. The synthesis and characterization of 2′,4,4′,6′-tetrahydroxychalcone in our lab was previously described (Wolf-Saxon et al. 2003). Michealis-Menten kinetics were performed at 25°C in 50mM HEPES-Na+ buffer pH 7.5 and 5% (v/v) EtOH as co-solvent, as previously established (Jez et al. 2000). *Mt*CHI-I was used for kinetics measurements at a final concentration of 40 nM. Time courses were monitored with a V-630 spectrophotometer (Jasco). To measure the effect of phloretin inhibition, phloretin was included with 0.5% DMSO as a co-solvent. Double reciprocal plots of inverse initial velocity *vs*. inverse substrate concentration were fitted using the linear fit function in Prism 10.0 (Graphpad). The value for K_i_ was calculated from each experiment using varying phloretin concentrations and reported as the mean with standard deviation. Three or more independent replicates were used to generate each datapoint.

## Author Contributions

Designed the research: J.R.B., M.A.R., E.R.W-S. Performed research: J.R.B., M.A.R., V.I.V.M., H.M.N., T.O.D., K.I., A.A.R., J.Z., E.R.W-S., C.C.M. Analyzed data: J.R.B., M.A.R., V.I.V.M., H.M.N., T.O.D., A.A.R., E.R.W-S. Writing, review and editing: J.R.B., M.A.R., V.I.V.M., C.C.M. All authors have read and approved the final manuscript.

## Acknowledgments

We thank the staff at the Advanced Light Source beamline 8.2.2, Lawrence Berkeley National Laboratory.

## Data availability Statement

Specific PDB information can be found at the Protein Data Bank for entry 38KF, Extended PDB ID pdb_000038KF.

## Abbreviations

CHI: chalcone isomerase
CHI-I: type-1 chalcone isomerase
CHI-2: type-2 chalcone isomerase
*Mt*CHI-I: *Medicago truncatula* type-1 CHI
DSF: differential scanning fluorimetry
App. K_d_: apparent equilibrium dissociation constant
ΔTm_max_: maximum thermal shift
RMS: root mean square

